# Uncoupling overeating and fat storage by modulation of different serotonergic receptors

**DOI:** 10.1101/2025.03.18.644037

**Authors:** Christina Clark, Rizaldy C Zapata, Ian R. Newman, Olivia Osborn, Michael Petrascheck

## Abstract

Psychotropic drugs such as antipsychotics improve symptoms of psychiatric disorders. However, they are associated with severe metabolic side effects that remodel energy balance, resulting in weight gain and increased food intake (hyperphagia). Here, we compare how antipsychotics and exogenous serotonin induce hyperphagia by remodeling energy balance. We find that the ability of serotonin and antipsychotics to remodel energy balance strictly depends on the serotonergic receptors SER-7 and SER-5, respectively. While both molecules induce hyperphagia, serotonin does so by increasing energy expenditure and reducing fat stores. In contrast, antipsychotics block the inhibitory effect of fat storage on feeding, thereby inducing hyperphagia and increasing fat stores. Thus, it is possible to manipulate energy balance to induce hyperphagia while either increasing or decreasing fat storage. Inactivation of the germline remodels energy balance similar to antipsychotic treatment, promoting hyperphagia while increasing fat storage. Consistent with overlapping mechanisms, antipsychotics are no longer able to remodel energy balance in both *C. elegans* and mice lacking an intact germline. Thus, our results uncouple overeating from fat storage and show that overeating can be induced by mechanisms that reduce or increase fat stores.

## Introduction

Over 900 FDA-approved drugs have been reported to cause metabolic side effects in humans, resulting in either weight loss or weight gain, by unknown mechanisms^1^. Stimulating drugs such as methylphenidate (Ritalin) can cause anorexia and weight loss^2–3^. Conversely, antipsychotics such as olanzapine are associated with weight gain and metabolic syndrome, at least in part caused by their tendency to stimulate overeating, called hyperphagia.^4–5^ Notably, patients on antipsychotics are more likely to develop diabetes than non-antipsychotic users of similar age, sex, and race^6^. Metabolic side effects are also the most commonly reported reason for patients to fail to comply with the antipsychotic regimen^7–8^. In many cases, it is unclear if the side effects are on or off-target and if the metabolic side effects are separable from the therapeutic effects.

Feeding is required for survival, and its regulation is at the core of maintaining metabolic health, reproduction, and aging. Feeding is a complex interaction between the various signaling pathways of the nervous system and the periphery and thus can only be studied *in vivo*^9–12^. From a basic science perspective, antipsychotics are an interesting experimental tool to modulate energy balance independently of the diet. The rapid onset of metabolic effects upon treatment with antipsychotics reveals that they must affect central mechanisms of energy balance controlled by energy intake, expenditure, and storage.

In *C. elegans*, feeding has been chiefly studied by measuring pharyngeal pumping, as defined by the rate of contractions of the pharynx, which has proven crucial to understanding feeding behavior^13–19^. Pharyngeal pumping does not, however, cover all aspects of feeding as it only measures the rate of pumping over a short period and does not capture the duration and frequency of feeding bouts. To remedy this, we developed the bacterial clearance assay, which measures the rate of decline of bacterial concentrations for defined populations of worms^8,20–21^. The bacterial clearance assay is comparable to the feeding studies conducted in mice and thus allows a better comparison of the effect of antipsychotics between the species^8,20,22–28^.

Here, we report that the administration of Antipsychotics, specifically loxapine and olanzapine, differentially modulate the energy balance of *C. elegans*. We show that antipsychotic-induced hyperphagia is distinct from serotonin-induced hyperphagia in several ways. Antipsychotics require a different G-protein coupled receptor (GPCR) and remodel different aspects of energy balance than serotonin. Antipsychotics induce hyperphagia by antagonizing the serotonergic GPCR SER-5, whereas serotonin induces hyperphagia by activating the serotonergic GPCR SER-7, as previously shown^8,20,29–30^.

Analyzing the effects of antipsychotics or serotonin on energy balance, we find that serotonin induces hyperphagia by increasing energy expenditure. In contrast, antipsychotics induce hyperphagia by raising the threshold at which fat storage inhibits feeding, leading to hyperphagia until a new ceiling in fat storage is reached. We show that the effect of antipsychotics on energy balance is similar to that of inactivating the germline. Inducing abundant fat storage by inactivating the germline also increases feeding and blocks the ability of antipsychotics to increase fat storage or hyperphagia further. In both cases, the increase in feeding is not explained by an increase in energy expenditure. The increase in feeding is associated with an increase in fat storage, suggesting that both interventions promote feeding by raising the threshold at which fat storage inhibits feeding. In contrast to antipsychotics, serotonin still induces hyperphagia in animals with an inactive germline, as it promotes hyperphagia by increasing energy expenditure and fat loss. We show the mechanism of antipsychotics to be evolutionarily conserved as ovariectomy in mice also increases weight gain and adiposity and blocks the ability of antipsychotics to promote it further.

## Results

### Antipsychotic and serotonin distinctly remodel energy balance

*C. elegans* feeding can be stimulated by either exogenous serotonin or treatment with antipsychotics, both inducing hyperphagia. Our previous findings showed that serotonin or antipsychotics induce hyperphagia via two genetically distinct receptors, *ser-7* and *ser-5,* respectively (**Fig. 1A-B**)^8^. For this study, we will distinguish between basal food intake, the food intake observed without stimulation, and hyperphagia, overeating observed when feeding is stimulated by either serotonin or an antipsychotic. Using the worm-based bacterial clearance assay^20^, which provides a highly accurate measure of food intake over time, we confirmed that hyperphagia induced by serotonin (5mM) strictly depends on the serotonin receptor *ser-7* but not *ser-5*. In contrast, overeating induced by the antipsychotic loxapine is dependent on the serotonin receptor *ser-5* and is blunted in *ser-7* mutants (**Fig. 1B-D**). Basal feeding was slightly but significantly increased in the *ser-5* knockout mutants *ser-5(vq1)*, which is consistent with the model in which loxapine antagonizes SER-5 signaling to induce feeding (**Fig. 1A, Sup1A**). As we will see below, the effect in the *ser-5(vq1)* knockouts is smaller than the effect caused by loxapine inhibition due to the different inhibition dynamics. We further confirmed the increase in feeding by measuring short-term feeding rates, counting pharyngeal pumps per minute (**Sup1B**)^18–19,21,31–36^. Thus, serotonin and loxapine both induce hyperphagia.

**Figure 1:**
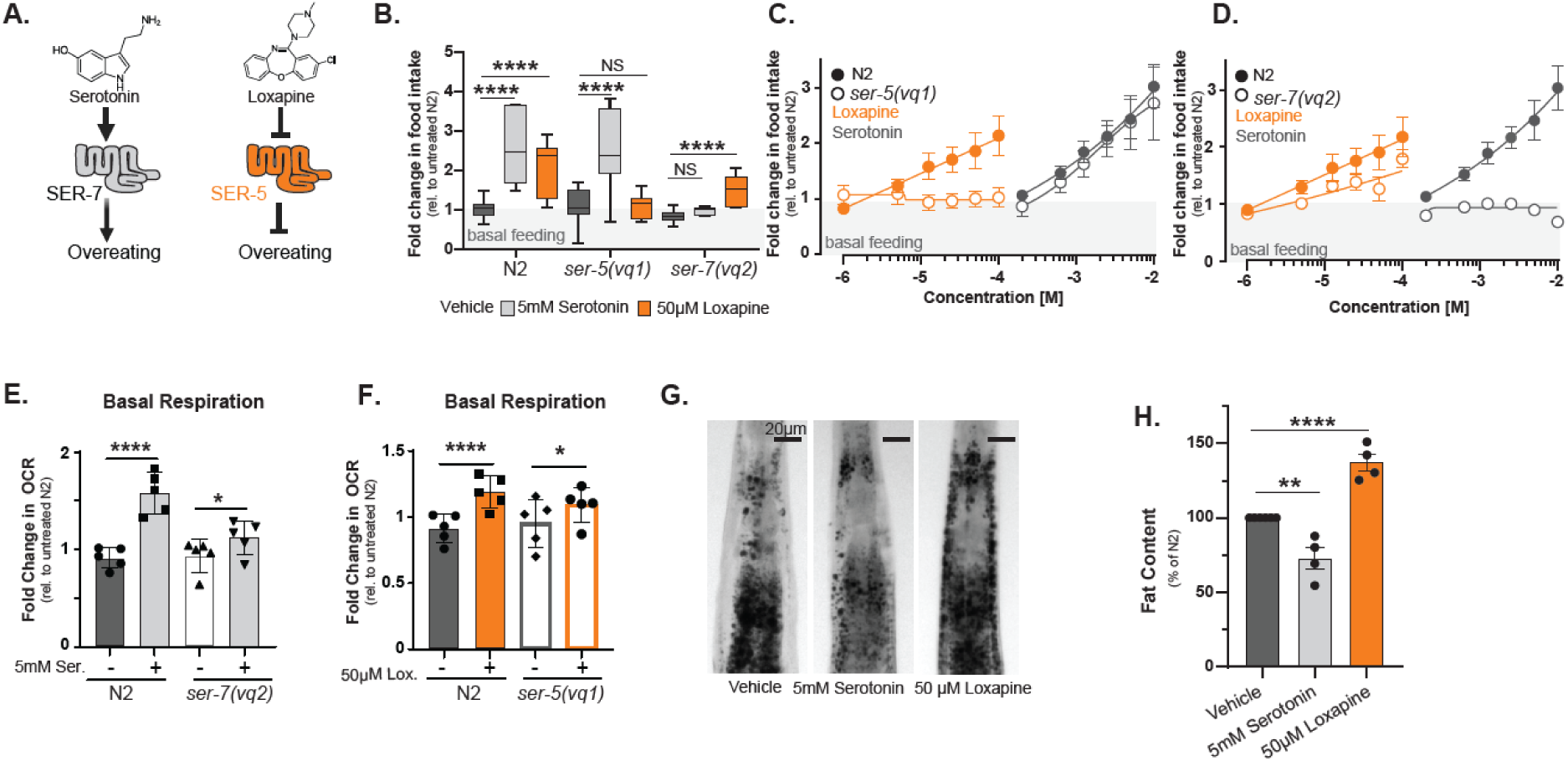
The antipsychotic, loxapine, and serotonin differentially remodel energy balance. **(A)** Model: Serotonin induces overeating by activating SER-7, while loxapine induces overeating by inhibiting SER-5. **(B)** Box blots representing the fold change in food intake of serotonin (light grey) and loxapine (orange) at optimal doses in N2 (wild-type), *ser-5(vq1)*, and *ser-7(vq2)* mutants. C-D Dose-response curves graphing fold change in food intake as a function of increasing serotonin and loxapine concentrations measured after 72 hours incubation on day 4 of adulthood. **(C)** Dose-response curves for N2 (wild-type) or *ser-5(vq1)* mutants treated with loxapine or serotonin, respectively **(D).** Dose-response curves for N2 (wild-type) or the serotonin mutant *ser-7(vq2)* treated with Loxapine or Serotonin, respectively. (**E-F**) shows bar graphs of basal oxygen consumption rates (OCR) measured on day 4 of adulthood, after 72h of treatment with either serotonin (5mM) or loxapine (50μM) measured on a Seahorse Extracellular Flux (XF) analyzer. OCR was normalized to the number of worms. Bars show the mean OCR of three trials with *n* = 100 animals each. **(E)** Oxygen consumption rate (OCR) of N2 or *ser-7(vq2)* mutants treated with vehicle (dark grey) or 5mM Serotonin (light grey) **(F).** Oxygen consumption rate (OCR) of N2 or *ser-5(vq1)* mutants treated with vehicle (dark grey) or 50 μM Loxapine (orange). **(G)** Representative images of vehicle, serotonin, or loxapine-treated adult *C. elegans* stained with Oil Red O to visualize fat depots. **(H).** Quantification of fat staining of day 4 adult N2 animals treated with either vehicle (dark grey), serotonin (light grey), or loxapine (orange). Bars show the mean % fat content relative to vehicle-treated controls of 4 trials consisting of n=20 animals each. All error bars indicate SEM. Images taken at 10X DIC. Asterisks indicate significance, \*\**p* < 0.05, \*\**p* < 0.01, \*\*\**p* < 0.001, *\*\*\*\*p*<0.00001 NS: not significant Abbreviation Ser. for serotonin and lox. for loxapine.

Next, we compared the impact of serotonin versus antipsychotic-induced remodeling of energy balance. Organismal energy balance is determined by energy intake, energy expenditure, and energy storage. To approximate energy expenditure, we measured oxygen consumption rates using a Seahorse XFe96 Flux Analyzer. Both serotonin and loxapine treatment increased oxygen consumption rates (OCR) (**Fig 1E-F, Sup 1 C-F**)^37–38^, dependent on their respective receptors, *ser-7* and *ser-5*. The increase in respiration was more pronounced for serotonin than for loxapine. However, in contrast to feeding, mutations in *ser-7* and *ser-5* severely blunted the increase in OCR but did not completely abolish it. This finding was consistent with previous findings that different serotonin receptors govern distinct aspects of energy balance^39^. Thus, serotonin and loxapine treatment increased energy expenditure via distinct receptors SER-7 and SER-5.

To compare the effects of serotonin or loxapine treatment on energy storage, we quantified fat depots by Oil Red O staining of animals treated with serotonin or loxapine for 72h starting on day 1 of adulthood^39^. Serotonin treatment of N2 adults decreased fat staining compared to age-matched vehicle controls, as previously reported ^35,39^, while loxapine treatment increased it (**Fig.1 G-H**).

Furthermore, serotonin failed to decrease fat staining in *ser-7(vq2)* knockouts, while loxapine increased it to the same extent as in N2 (**supp Fig, 1G, H**). In untreated *ser-5(vq1)* knockouts, fat staining was increased to the same extent as observed in loxapine-treated N2 animals(**supp Fig, 1G, H)**. Loxapine treatment no longer increased fat staining in *ser-5(vq1)* knockouts but reduced it to wild-type levels (**Sup 1H**). Thus, serotonin and loxapine treatment have opposing effects on energy storage, mediated by their respective receptors SER-7 and SER-5.

Taken together, these results show that serotonin induces hyperphagia and energy expenditure (OCR) by signaling via SER-7 and reduces fat storage. More surprising was the effect of loxapine on energy balance. Loxapine also induces hyperphagia, slightly raises energy expenditure, and, in contrast to serotonin, substantially increases fat storage (**Fig. 1E-G**). They suggest that SER-5 directly or indirectly mediates the inhibition of feeding by fat stores. Inhibiting SER-5 by adding loxapine to adult animals suddenly blocks fat-mediated inhibition of feeding, resulting in the induction of hyperphagia until fat stores reach a maximum level (**Fig. 1G, supp Fig. 1 G, H**). Since the *ser-5(vq1)* knockout lacks SER-5 signaling throughout development by the time it reaches adulthood, fat storage has already reached the higher threshold necessary to inhibit feeding, resulting in a much smaller hyperphagic effect (**supp Fig. 1 G, H**). This gain in adiposity by treatment with antipsychotics is similar to what is observed in mice, where the antipsychotic-mediated disruption of leptin signals shifts the ratio of fat stores to feeding^40^.

### Antipsychotics and serotonin induce overlapping transcriptional signatures through distinct serotonin receptors

To better understand the molecular basis of serotonin or loxapine treatment on energy balance, we set out to identify differentially expressed genes (DEGs) induced by either treatment. Because each molecule remodeled the energy balance by a specific receptor, we determined how serotonin and loxapine treatment changed gene expression in wild-type N2 animals and how they changed gene expression in *ser-7(vq2)* and *ser-5(vq1)* mutants.

For each sample collected for RNAseq, we confirmed the induction of hyperphagia by serotonin or loxapine treatment. MDS plots revealed that the induction of hyperphagia separated gene expression patterns between all samples, more so than genetic background or pharmacology. Serotonin or loxapine-treated N2 animals that overate formed one cluster. In contrast, all other samples that did not overeat, either because they were vehicle-treated or because mutations in *ser-7* or *ser-5* prevented hyperphagia induction, formed a separate cluster (**Fig. 2A**). Thus, the effect of hyperphagia on gene expression dominated the effects of genotype or pharmacology. Consistently, there were only minor gene expression differences between N2 and *ser-7(vq2)* vehicle-treated control animals (257 DEGs) and N2 and *ser-5(vq1)* vehicle-treated control animals (5 DEGs). As expected, the most differentially expressed genes were the mRNAs for *ser-7* and *ser-5*, respectively (**Sup Fig 2A/B**).

**Figure 2:**
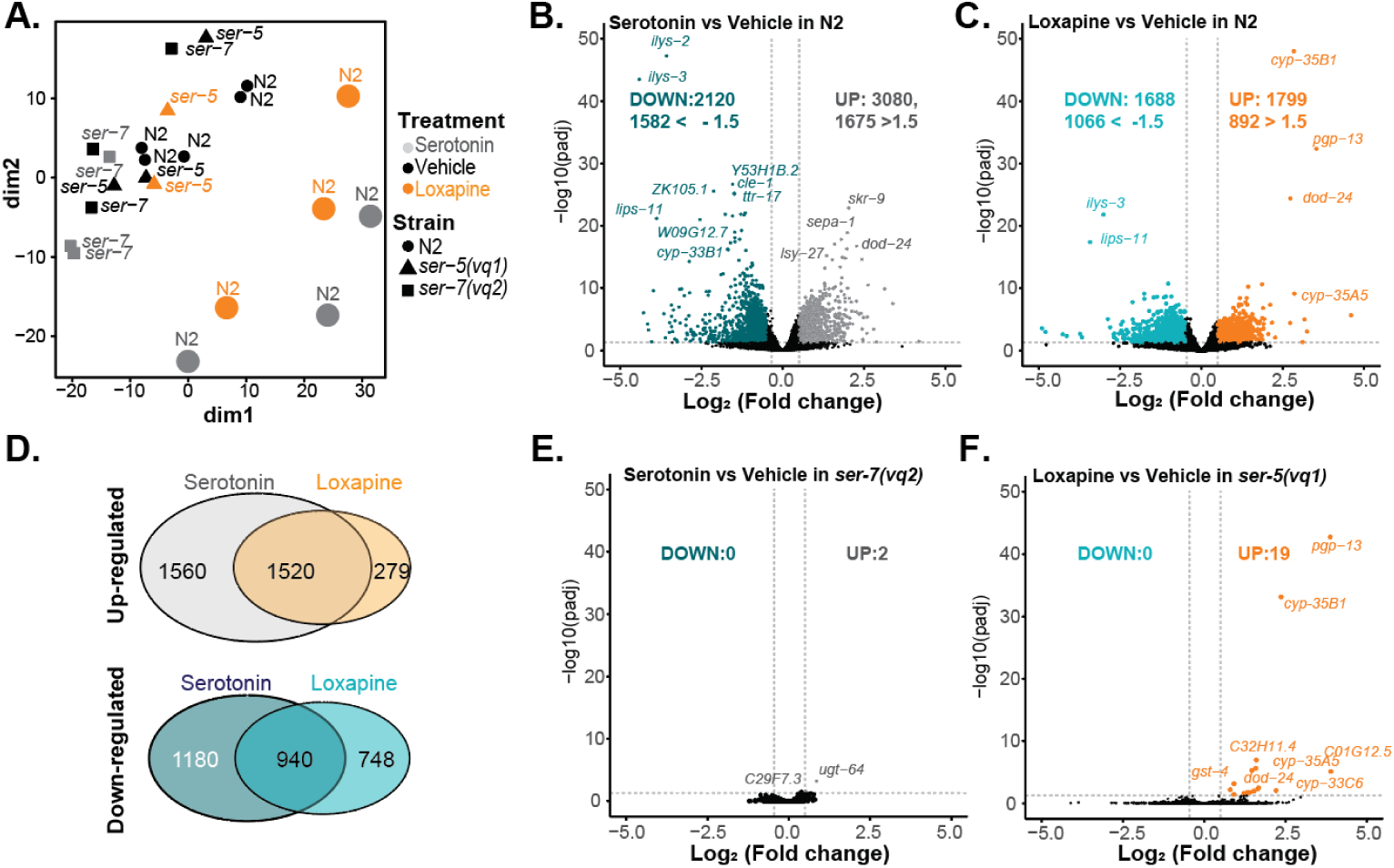
SER-5 and SER-7 are the sole receptors mediating the transcriptional response to hyperphagia. **(A)** Multidimensional scaling (MDS) plot of the RNAseq samples. Samples of overeating animals (grey and orange circles) cluster separately from normally eating animals. B-C and E-F show volcano plots representing -Log_10_ (pvalues) as a function of Log_2_(Fold changes) in gene expression for 14,912 genes detected in all samples. Differentially expressed genes (DEGs) are colored. **(B)** Serotonin treatment-induced gene expression changes in N2 animals **(C).** Loxapine treatment-induced gene expression in N2 animals. **(D).** Venn diagram of DEGs showing the overlap of up-regulated (top) and down-regulated DEGs from serotonin or loxapine treated samples depicted in **B, C**. **(E).** Serotonin treatment-induced gene expression changes in *ser-7(vq2)* mutants **(F).** Loxapine treatment-induced gene expression changes in *ser-5(vq1)*. For all volcano plots FDR ≥ 0.05.

Treatment of N2 animals with either serotonin or loxapine resulted in dramatic gene expression changes. Serotonin treatment induced 5200 DEGs, of which 3080 were up, and 2120 were downregulated (adjusted P <0.05) (**Fig. 2B**). Loxapine treatment of N2 animals resulted in 3487 DEGs, of which 1799 were up, and 1688 were downregulated (adjusted p <0.05) (**Fig. 2C**). As hinted by the MDS plot, both serotonin and loxapine induced overlapping transcriptional signature with a correlation factor of R= 0.66 (P<10^-15^) for the whole dataset and a correlation factor of R= 0.94 for the 2460 shared DEGs. Among the 2460 shared DEGs, 1520 DEGs were up-regulated by both treatments (**Fig 2D**). These 1520 up-regulated DEGs represented 84% of all DEGs up-regulated by loxapine, suggesting that the loxapine response is a subset of the serotonin response. The interpretation that the transcriptional response to loxapine represents a subset of the response to serotonin was also reflected in the MDS plot, where the loxapine-treated N2 samples are found between the control and serotonin-treated samples (**Fig. 2A)**

The transcriptional responses to both molecules were completely dependent on their respective receptors. Serotonin treatment of *ser-7(vq2)* mutants resulted in 2 DEGs (**Fig. 2E**). Loxapine treatment of *ser-5(vq1)* mutants resulted in 19 DEGs, almost all of which are classical xenobiotic response genes, including cytochromes (cyp) or ABC transporters (pgp). Over 99.4% of the transcriptional responses to serotonin or loxapine treatment observed in wild-type N2 animals were abolished in *ser-7(vq2)* or *ser-5(vq1)* mutants, respectively. These results show that SER-7 and SER-5 are the main and possibly only targets for serotonin and loxapine to remodel transcription.

### Transcription and translation of a shared transcriptional signature are necessary to induce hyperphagia

The MDS plot and the considerable overlap of 2460 shared DEGs revealed that serotonin and loxapine induce a shared transcriptional signature (**Fig. 2D**). As this shared transcriptional signature was induced by different receptors, we reasoned that it is most likely related to the shared ability of both molecules to induce hyperphagia (**Fig. 2B-F**). We analyzed the shared transcriptional signature of 1520 up-regulated DEGs for gene set enrichment using WormCat 2.0^41^. The four highest-ranked terms were DNA, mRNA functions, proteolysis by the proteasome, and metabolism (**Fig 3A, B**)^41^. These GO terms suggested a transcriptional response that promotes the expression of genes that direct the synthesis of endogenous biomolecules to process the ingested nutrients. Furthermore, the knockdown phenotypes of these 1520 DEGs were enriched for phenotypes like embryonic lethality (P= 8.5*10^-51^), sterile progeny (P= 1.23*10^-14^), and reduced brood size (P= 1.6*10^-12^) as determined by EnrichR ^42–43^.

**Figure 3:**
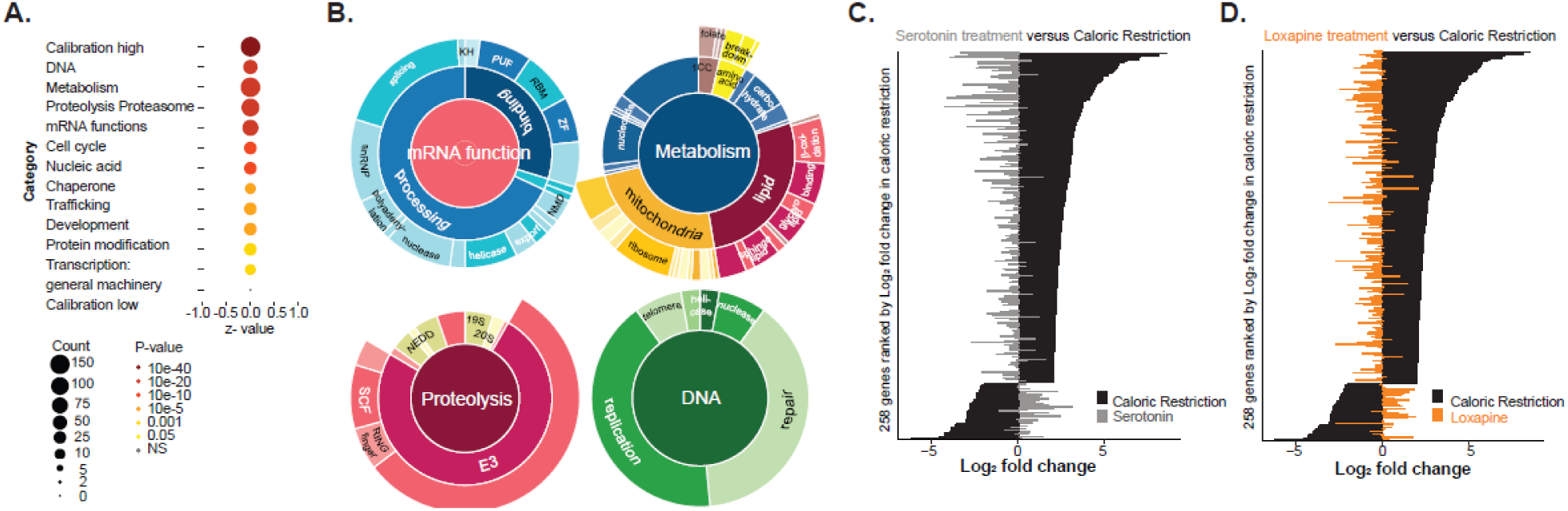
Biosynthetic pathways up-regulated by serotonin or loxapine treatment. The 1520 DEGs shared between serotonin or loxapine-treated samples (Fig. 2D) were analyzed by gene set enrichment. **(A)** Gene Ontology analysis by WormCat 2.0 of 1520 shared DEGs. **(B)** Sunburst graphs represent the four highest-ranked Gene Ontologies and their respective subcategories enriched in the 1520 DEGs shared in serotonin and loxapine-treated animals. Graphs in **C-D** show bar graphs depicting the Fold changes in gene expression of 258 genes in fasting animals subjected to caloric restriction or overeating animals treated with either serotonin (**C**) or Loxapine (**D**). CR-Data was taken from PMID 23935515. **(C)** Hyperphagia induced by serotonin (grey) shows opposing gene expression patterns to fasting-induced by caloric restriction (black). **(D)** Hyperphagia induced by loxapine (orange) shows opposing gene expression patterns to fasting-induced by caloric restriction (black).

We further explored the idea that the observed transcriptional changes were induced to process the ingested nutrients. Provided that the purpose of shared transcriptional changes was to process nutrients ingested by feeding, the direction of their expression should change in the opposite direction upon fasting. We compared our hyperphagic transcriptomes to transcriptomes of *C. elegans* subjected to calorie restriction (CR)^44^. First, we selected DEGs whose expression level changed by a factor of 4 or more in either direction under conditions of CR (258 DEGs) published by Heestand BN et al. ^44^. We plotted their Log2 fold expression changes in response to CR and serotonin (**Fig. 3C**) or CR and Loxapine treatment (**Fig. 3D**). The plots in **Fig. 3** show that the expression of genes responding to fasting (CR) inverts upon the induction of overeating by serotonin or loxapine treatment. This inverted expression pattern was not restricted to a few genes (**Fig.3C, D** but held for 2148 significantly changed DEGs that were represented in all three datasets. Correlating the expression of DEGs identified for CR resulted in strong negative correlations between fasting and overeating (R _serotonin_= -0.41, P= 1.6*10^-88^; R _Loxapine_ = -0.27; P= 7.4*10^-38^).^44^ Consistent with the interpretation that the observed gene expression changes are directly related to the processing of ingested nutrients, the hyperphagic effect of serotonin treatment is greater than that of loxapine and shows a stronger negative correlation. Thus, we concluded that the shared transcriptional signature induced in serotonin or loxapine-treated animals is a response to ingested nutrients that are necessary to process them.

Next, we set out to test if the shared transcriptional signature is necessary to induce hyperphagia. We asked if blocking of transcription, mitochondrial respiration, or proteasomal degradation by chemical inhibitors would block hyperphagia induced by serotonin or loxapine. We co-treated vehicle, loxapine, or serotonin-treated animals with increasing doses of (i) the RNA polymerase inhibitor actinomycin D to reduce transcription, (ii) the complex III inhibitor antimycin A to reduce mitochondrial respiration, (iii) the proteasome inhibitor, bortezomib to reduce protein degradation and (iv) the translation elongation inhibitor anisomycin to lower protein synthesis. The quantification of food intake revealed that all of these inhibitors either abolished or at least dampened hyperphagia, as well as basal feeding (**Fig. 4A-D**).

**Figure 4:**
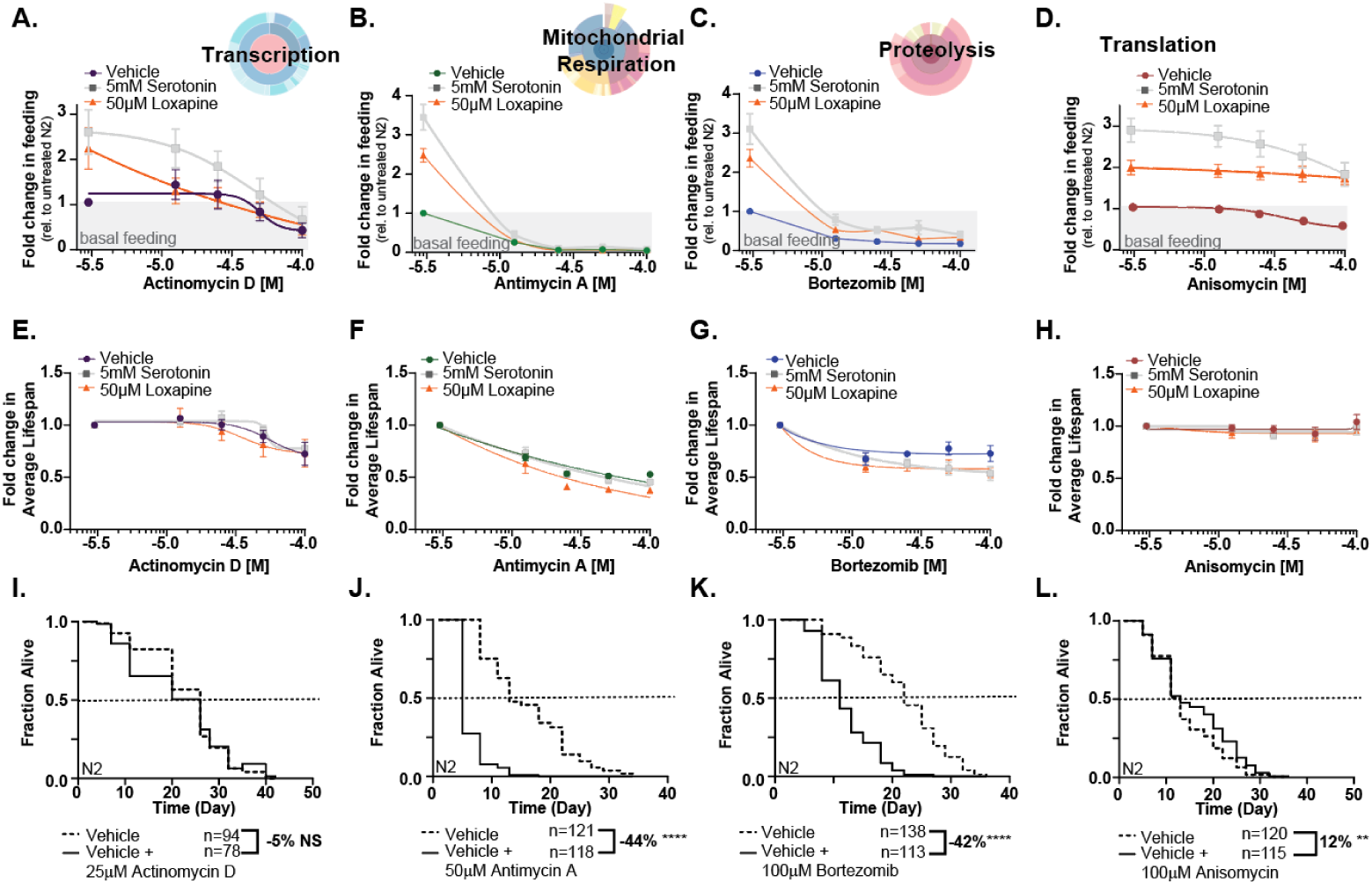
Chemical inhibition of biosynthetic pathways suppresses loxapine and serotonin-induced hyperphagia. Graphs A-D show a fold change in food intake relative to the N2 vehicle control as a function of increasing inhibitor concentration of vehicle, serotonin (5mM), or loxapine (50uM) treated N2 wild-type animals. The biological process that is inhibited is indicated on top. **(A)** Dose-response curve depicting the fold change in feeding as a function of increasing doses of transcription inhibitor Actinomycin D. **(B)** Same as (A) but for the mitochondrial respiration inhibitor Antimycin A. **(C)** Same as (A) but for the proteasome inhibitor bortezomib. **(D)** Same as (A) but for the translation inhibitor anisomycin. Graphs E-H show the average fold change in lifespan for N2 worms as a function of increasing doses of the inhibitor used in A-D, alone or in combination with serotonin ( 5mM, grey) or loxapine (50μM, orange). **(E)** Dose-response curves depicting the fold change in lifespan as a function of increasing doses of Actinomycin D. **(F)** Same as (E) but for Antimycin A. **(G)** Same as (E) but for Bortezomib. **(H).** Same as (E) but for Anisomycin. Graphs I-L show Kaplan-Meier survival curves for N2 treated with the same inhibitors as in A-D. **(I)** Kaplan-Meier survival curves for N2 treated with Actinomycin D. **(J)** Kaplan-Meier survival curves for N2 treated with Antimycin A. **(K)** Kaplan-Meier survival curves for N2 treated with bortezomib. **(L)** Kaplan-Meier survival curves for N2 treated with anisomycin. Survival data were analyzed by a Log-rank test. Asterisks indicate significance, \*\**p* < 0.05, \*\**p* < 0.01, \*\*\**p* < 0.001, *\*\*\*\*p*<0.00001 NS: not significant Abbreviation Ser. for serotonin and lox. for loxapine.

However, antimycin A and bortezomib treatment appeared to be toxic. Conducting lifespan assays for all inhibitors confirmed that antimycin A and bortezomib were toxic and shortened lifespans (**Fig. 4E-H**). For the transcription inhibitor actinomycin D we were able to identify concentrations that did not shorten lifespan yet abolished hyperphagia. The translation inhibitor anisomycin was not toxic at any concentrations, yet dampened feeding (**Fig. 4D, H, L**, **supp Fig.4**). The dampening effect of anisomycin on loxapine-induced hyperphagia was mild and only became apparent after normalization (**supp Fig. 4D**).

Overall, these results support the hypothesis that the shared transcriptional response induced by serotonin and loxapine is not a direct result of modulating receptor signaling but a necessary secondary response to hyperphagia. To maintain feeding, ingested nutrients need transformation into endogenous biomolecules. Blocking transformation into endogenous biomolecules abrogates feeding. The key result that places the transcriptional response downstream of hyperphagia is that *ser-5(vq1)* knockouts show increased fat storage but very limited hyperphagia and almost no change in transcription. Thus, the observed changes in transcription can neither be the cause nor the direct consequence of the accumulation of fat.

### Increased fat storage prevents antipsychotic-induced but not serotonin-induced hyperphagia

The inhibitor experiments revealed that inhibition of translation reduces hyperphagia without being overtly toxic. This observation allowed us to investigate potential tissue-specific contributions to hyperphagia. To tissue-specifically block translation, we used the auxin (indole-3-acetic acid, IAA) inducible degron (AID) system. The AID system enables temporal and tissue-specific degradation of proteins. We chose to degrade RNA polymerase I, called POLI in mammals or *rpoa-2* in *C. elegans.* POLI/RPOA-2 transcribes rRNA in the nucleolus and thus controls ribosomal biogenesis^45–47^. In the AID system, the protein of interest (POLI/*RPOA-2*) is tagged with a degron that interacts with TIR1 upon the addition of IAA. TIR1, a plant protein, recruits ubiquitin ligases upon binding to IAA, which then degrade the degron-tagged protein. The necessity of IAA to induce degradation provides a temporal control, while the necessity of TIR1, which can be expressed tissue-specifically, enables spatial-/tissue-specific control. Thus, the auxin-induced degradation of POLI/RPOA-2 will block rRNA transcription and thus prevent ribosome biogenesis, severely limiting translation.

We first asked how limiting translation in different tissues would affect basal feeding. As expected, limiting translation by degradation of POLI/RPOA-2 in the whole body dramatically reduced feeding (**Fig. 5A**, *eft-3p*::TIR1) and the animal’s size (**Fig. 5B**). Limiting translation in specific tissues, such as the hypodermis, pharynx, muscle, and intestine also reduced feeding (**Fig. 5A**, *col-10, myo-2, myo-3, ges-1*) and body reduced size, except for limiting translation in the pharynx (*myo-2*), which did not affect body size significantly (**Fig. 5B**). The hypodermis and the intestine are the largest organs in *C. elegans*, each representing about ∼25% of the animal in volume, followed by the germline and body wall muscles representing ∼18%^48^. The tissue that caused the most significant reduction in feeding upon limitation of translation was the hypodermis, followed by the intestine and body wall muscles^49–52^.

**Figure 5.**
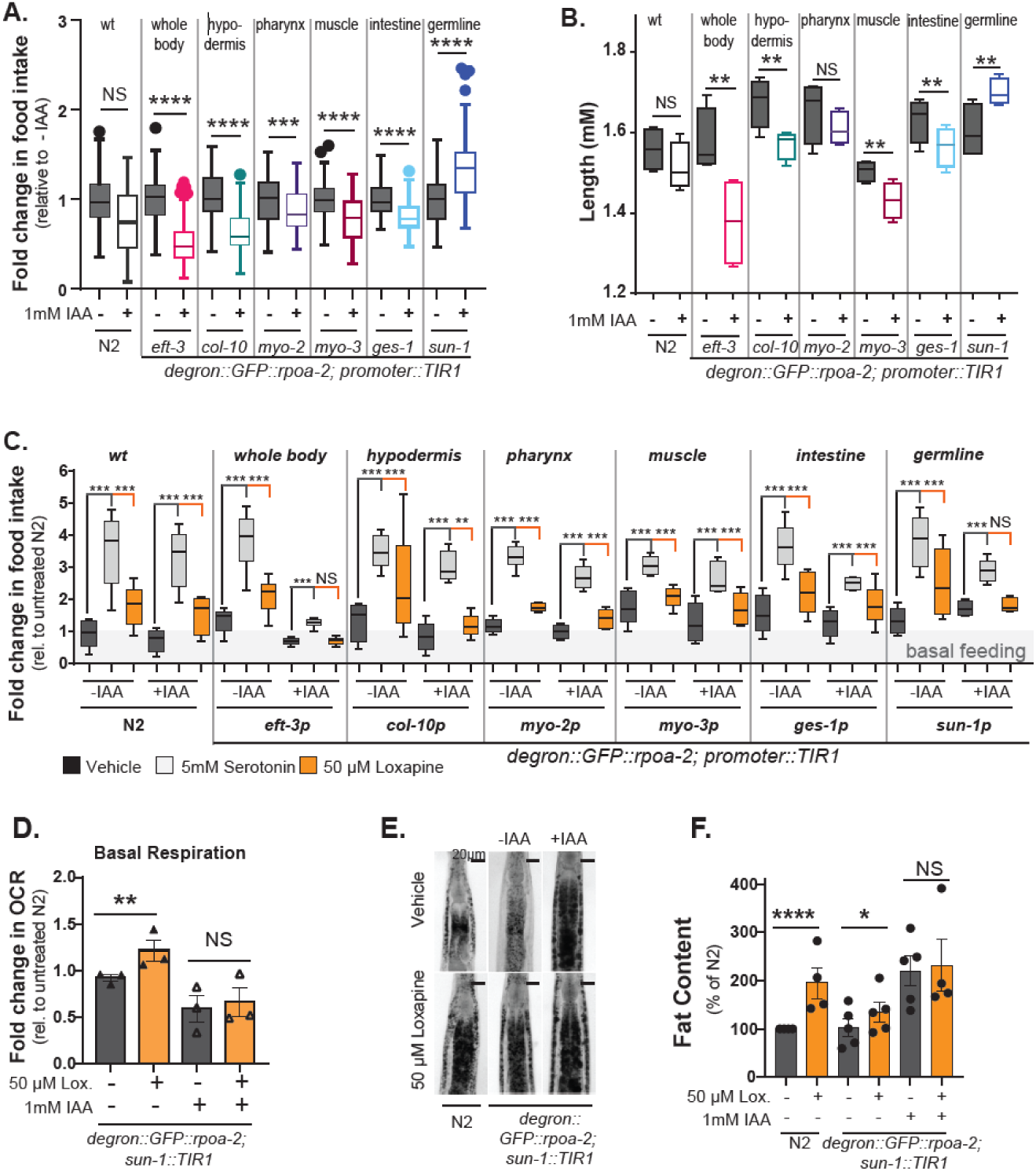
Translation in the germline is required for antipsychotic-induced overeating. Tissue-specific inhibition of translation is achieved by fusing RNA polymerase I (*rpoa-2*) to a degron (*degron*::*GFP*::*rpoa-2*) and by the tissue-specific expression of the plant protein TIR1, required for degradation induced by the addition of IAA. TIR1 expression is driven by the following promoters: whole body (*eft-3p*), epidermis (*col-10p*), pharynx *(myo-2p*), body wall muscle (*myo-3p*), intestine *(ges-1p*), and germline (*sun-1p*). To induce RPOA-2 degradation, the transgenic strains were treated with 1 mM IAA for 72h. **(A)** Box plots of fold change in food intake of N2 (wild-type) animals or *degron*::*GFP*::*rpoa-2* transgenic animals treated with vehicle or 1mM IAA to limit translation by inducing the degradation of RPOA-2 in the tissues indicated at the top. **(B)** The average length of N2 (wild-type) animals or *degron*::*GFP*::*rpoa-2* transgenic animals treated with vehicle or 1mM IAA to limit translation by inducing the degradation of RPOA-2 in the tissues indicated at the top. **(C)** Box plots of fold change in food intake in response to vehicle (black), serotonin (grey, 5mM), or loxapine (orange 50 μM) treatment of N2 (wild-type) animals or *degron*::*GFP*::*rpoa-2* transgenic animals treated with vehicle or 1mM IAA to limit translation by inducing the degradation of RPOA-2 in the tissues indicated at the top. **(D)** Fold change in oxygen consumption rate (OCR) relative to vehicle control animals of *degron*::*GFP*::*rpoa-2* transgenic animals treated with vehicle (dark grey) or 50 μM Loxapine (orange) in which germline translation remained intact (-IAA) or was blocked by the IAA induced degradation of ROPA-2 (+IAA). Each dot represents a trial of *n* = 100 animals. **(E).** Representative images of Oil Red O staining of vehicle or loxapine (50 μM) treated N2 or *degron*::*GFP*::*rpoa-2* transgenic animals in which the IAA-induced degradation of ROPA-2 limited germline translation. **(F).** Quantification of the percent fat content in vehicle-treated or Loxapine-treated (orange) N2 or *degron*::*GFP*::*rpoa-2* transgenic in which the IAA-induced degradation of ROPA-2 limited germline translation. Each dot in F represents a trial consisting of n=30 animals. All error bars = SEM. Images taken at 10X DIC. Asterisks indicate significance, \*\**p* < 0.05, \*\**p* < 0.01, \*\*\**p* < 0.001, *\*\*\*\*p*<0.00001 NS: not significant Abbreviation Ser. for serotonin and lox. for loxapine.

Surprisingly, the degradation of POLI/RPOA-2 in the germline (*sun-1p*::TIR1) increased food intake and body size (**Fig. 5A, B**). The protein synthesis activity in the germline must be considerable to account for the number of eggs produced. Thus, we expected that amino acids ingested by feeding would be used in the germline to promote protein synthesis and that limiting germline translation would reduce feeding. We tested how severely the degradation of POLI/RPOA-2 in the germline would affect its reproductive function and found that it rendered the animals completely sterile. Thus, limiting translation in the germline blocks reproduction but promotes hyperphagia. These data suggest that inactivation of the germline restructures energy balance, resulting in increased feeding and somatic growth (**Fig. 5A, B,** *sun-1p*::TIR1).

Limiting translation by the degradation of POLI/RPOA-2 in any tissue tested did not block the ability of serotonin to induce hyperphagia. The hyperphagic effect of serotonin was blunted by limiting translation in the whole body (**Fig. 5C**, *eft-3p*::TIR1) and to a lesser degree in the intestine (**Fig. 5C**, *ges-1p*::TIR1). However, even when translation was limited in the whole body, serotonin treatment increased food intake compared to vehicle-treated control animals. In contrast, degradation of POLI/RPOA-2 in the whole body (**Fig. 5C**, *eft-3p*::TIR1) or in the germline (**Fig. 5C**, *sun-1p*::TIR1) blocked loxapine-induced hyperphagia.

The finding that limiting translation in the germline blocked loxapine-induced hyperphagia raised the question of whether the block was due to inhibiting translation or due to the lack of a germline (**Fig. 5C**). However, the comparatively mild effect of anisomycin inhibiting loxapine-induced hyperphagia pointed to a germline effect (**Fig. 4D**). In agreement with this view, previous studies revealed increased fat storage in germline-less *glp-1(ts)* animals suggesting remodeling of energy balance in animals lacking an active germline^53^. Measuring energy expenditure and storage revealed that limiting translation in the germline reduced energy expenditure, as measured by oxygen consumption rates, by almost ∼50%. It further increased energy storage by doubling the amount of fat stored (**Fig. 5D-F**), as previously observed in *glp-1(ts)* germline-less mutants^53^. Both the remodeling of energy balance by inactivating the germline and treatment with loxapine were accompanied by an increase in fat storage, revealing that both mechanisms perturb the balance between fat storage and feeding.

Inactivating the germline by limiting translation also blocks loxapine from remodeling any aspect of energy balance. In the absence of a functional germline, loxapine failed to increase feeding or fat accumulation further beyond the already elevated levels (**Fig. 5E, F**). However, serotonin was still able to increase feeding in the absence of a functional germline (**Fig. 5C**), showing that the animal’s metabolism remained responsive to other hyperphagic induction mechanisms. Because loxapine treatment and the lack of a functional germline both increase feeding and fat accumulation, it is not clear if they are part of the same pathway or a parallel pathway with the same outcome. The most parsimonious explanation is that both interventions suddenly raise the threshold at which fat storage inhibits feeding. As a consequence, the animals start to overeat until the fat storage reaches a new threshold, at which point the overeating fades out. Thus, loxapine no longer promotes hyperphagia beyond what is seen in germline-less animals as the higher fat storage threshold is already reached.

### Increasing fat storage by ovariectomy blocks antipsychotic-induced weight gain in mice

The antipsychotic best known to disrupt human energy balance is olanzapine^54–56^. Olanzapine increases hyperphagia in wild-type N2 animals, similar to loxapine. Olanzapine also increased hyperphagia in the *sun-1p*::TIR1 transgenic strain without IAA treatment (**Fig. 6A**). However, upon treatment with IAA to induce the degradation of POLI/RPOA-2, olanzapine treatment no longer induced hyperphagia beyond the increased level of feeding seen due to the impaired germline (**Fig. 6A**). Similarly, olanzapine increased fat accumulation in wild type N2 but not in *sun-1p*::TIR1 transgenics without germline in which the fat storage was already abundant (**Fig. 6B, C**). Finally, olanzapine did not increase the oxygen consumption of IAA-treated *sun-1p*::TIR1 transgenics lacking an active germline (**Fig. 6D**). Thus, all our findings for loxapine extend to other antipsychotics, such as olanzapine.

**Figure 6:**
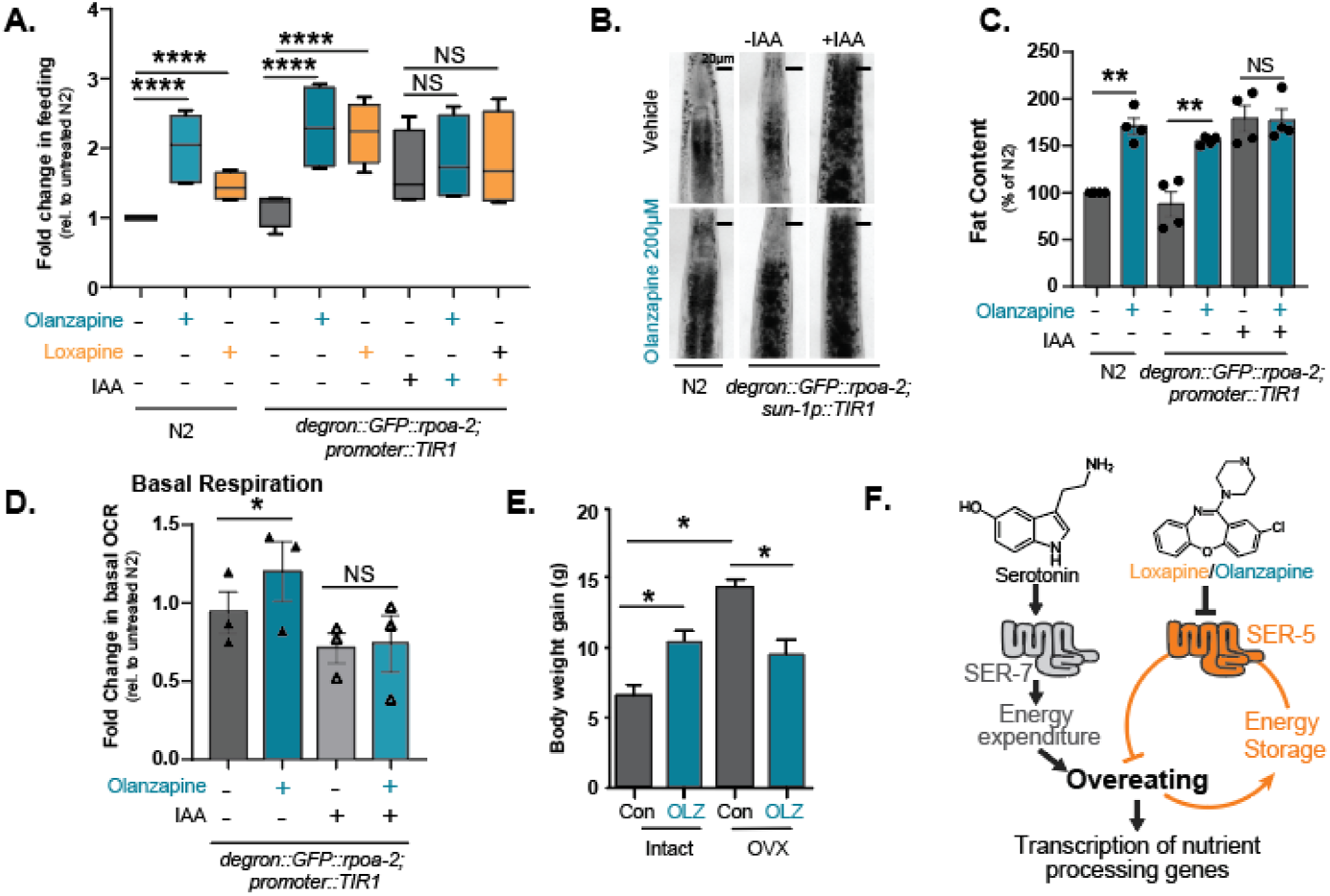
Germline Control of Antipsychotic Induced Overeating. **(A).** Fold Change in Food intake for N2 (wild-type) and *degron::GFP::rpoa-2* (germline) transgenic animals treated with loxapine (50 μM, orange) and olanzapine (200 μM, teal). The addition of IAA induces the degradation of RPOA-2 in the germline to limit translation. **(B)** Representative images of Oil Red O staining to quantify fat storage of N2 animals *degron::GFP::rpoa-2* (germline) transgenic animals treated with vehicle or 200 50 μM Olanzapine. Images taken at 10X DIC. **(C)** Quantification of the percent fat content in vehicle-treated (dark grey) or olanzapine-treated (teal) animals with intact germline (IAA) or animals lacking RPOA-2 germline expression. Each dot represents a trial consisting of n=40 animals. **(D)** OCR of vehicle or olanzapine (200 μM) treated *degron::GFP::rpoa-2* transgenic animals with an active (-IAA) or inactive germline (+ IAA) due to the degradation of RPOA-2. Measurements were taken on a Seahorse Extracellular Flux (XF) analyzer. Each dot represents a trial of n = 100 animals. **(E).** Change in body weight of female C57BL6 mice given olanzapine (54mg/ml in the chow) with or without an ovariectomy. **(F).** Model of how loxapine and serotonin remodel energy balance. Asterisks indicate significance, \*\**p* < 0.05, \*\**p* < 0.01, \*\*\**p* < 0.001, *\*\*\*\*p*<0.00001 NS: not significant Abbreviation Ser. for serotonin and lox. for loxapine.

The increase in fat accumulation and decrease in respiration in response to an inactive germline was reminiscent of the effects of ovariectomy in mice^57^. Ovariectomy in mice increases body weight and reduces oxygen consumption, similar to the effects we observed in *C. elegans* upon inactivation of the germline. We, therefore, asked if the metabolic side effects of olanzapine in mice are also dependent on an intact germline. We thus fed female C57BL6 mice that were either ovariectomized or sham-treated and were either fed control chow or chow-containing olanzapine (54mg/kg). After 4 weeks, we measured their body weight. As previously established, olanzapine treatment and ovariectomy both increased body weight^50^. However, as in *C. elegans*, olanzapine was unable to increase the already elevated body weight in ovariectomized mice and even decreased it. Thus, the metabolic side effects of antipsychotics are also blocked in mice lacking an active germline (**Fig. 6E**), which were previously shown to have increased adiposity and lower oxygen consumption^57^, similar to the effects seen in *C. elegans*.

## Discussion

Psychotropic drugs such as antipsychotics remodel energy balance in humans by mechanisms that are poorly understood. Here, we investigated how antipsychotics such as loxapine or olanzapine remodel energy balance in *C. elegans*. We compared their effects to those of serotonin, the only other known inducer of hyperphagia in *C. elegans*. We find that serotonin and antipsychotics both increase hyperphagia by acting through the serotonergic receptors SER-7 and SER-5, respectively. The main difference in remodeling energy balance between the two molecules was in their effect on fat storage. While serotonin treatment reduces fat storage, antipsychotics increase it (**Fig. 1**, **6, B, C, F**), consistent with its side effects seen in humans^1,4–5^. Evaluating the different aspects of energy balance revealed that serotonin drives hyperphagia by increasing energy expenditure via SER-7 signaling, thereby lowering fat storage. In contrast, loxapine drives hyperphagia by inhibiting SER-5, which directly or indirectly mediates the inhibition of feeding by fat stores. Inhibiting SER-5 raises the threshold at which stored fat starts to inhibit feeding, resulting in hyperphagia until the higher fat storage threshold is reached (**Fig. 6F**). The results show that it is possible to uncouple overeating from fat storage and to manipulate *C. elegans* energy balance to induce hyperphagia while either increasing or decreasing fat storage.

The model presented in **Fig. 6F** describing how serotonin remodels energy balance is supported by the following results. Serotonin induces hyperphagia (**Fig. 1B-D**), increases respiration (**Fig. 1E**), and decreases fat storage (**Fig. 1H, G**). The decrease in fat storage is the result of the large increase in respiration, whose need for energy is not sufficiently met by the increase in feeding, thus depleting fat stores. Serotonin remodels all aspects of energy balance via signaling by the serotonin receptor SER-7, a GPCR that couples to Gαs. Gαs activates adenylate cyclase, which then stimulates lipolysis^58–59^, consistent with the model in **Fig. 6F**.

The model presented in **Fig. 6F** describing how antipsychotics remodel energy balance is supported by the following results. Loxapine induces hyperphagia (**Fig. 1B-D**), slightly increases respiration (**Fig. 1F**), and substantially increases fat storage (**Fig. 1H, G**). These effects were largely independent of SER-7 but required SER-5^8^. Increased fat storage tends to inhibit feeding or is associated with lower respiration, raising the question of how loxapine treatment remodeled energy balance. Heterologous expression of SER-5 in HEK293T cells showed that SER-5 is a serotonin receptor that responds to serotonin and that it might be coupled to arrestin, suggesting an inhibitory role^60^. Consistent with an inhibitory role in feeding, inhibiting SER-5 by loxapine induces hyperphagia. In addition, *ser-5(vq1)* knockout animals show a small but significant increase in feeding compared to wild-type animals (**Fig. 1B, D, supp Fig. 1A**). Both loxapine-treated or *ser-5(vq1)* knockout animals show a substantial increase in fat storage (**Fig. 1H, G**).

The combination of these observations is consistent with a model in which SER-5 mediates the inhibitory effect of stored fat on feeding. Inhibiting SER-5 by adding loxapine to adult animals suddenly releases the inhibition of fat on feeding. The resulting disinhibition of feeding leads to hyperphagia until fat storage reaches its maximal capacity (**Fig. 1G, supp Fig. 1 G, H**). The much smaller hyperphagia in the genetic *ser-5(vq1)* knockouts compared to loxapine treatment is the result of a permanent lack of SER-5 signaling throughout development. By the time *ser-5(vq1)* knockouts reach adulthood, their fat storage has already reached maximal fat storage, limiting hyperphagia (**supp Fig. 1 G, H**). The final piece of evidence that increasing the threshold at which fat storage inhibits feeding can promote hyperphagia came from the experiments inactivating the germline (**Fig. 5**). Inactivating the germline by limiting translation increased fat storage and promoted feeding while reducing respiration. Antipsychotics were no longer able to alter any of the metabolic parameters in animals lacking a functional germline, both in *C. elegans* and mice (**Fig. 5**, **6**). The interpretation that SER-5 acts as a “stop eating” receptor when activated by serotonin is also consistent with findings that SER-5 is involved in the adaptive avoidance of CuSO4, a toxin that, when ingested, would harm the animal^61^.

The most striking demonstration that SER-7 and SER-5 were the main targets of serotonin and loxapine came from the analysis of gene expression. While treatment with both molecules induced thousands of transcriptional changes, over 99% of these changes did not occur in the respective mutants (**Fig. 2**). The initial goal when conducting the gene expression analysis was to understand the mechanisms by which SER-5 and SER-7 signaling modulated energy balance. However, the *ser-7(vq2)* and *ser-5(vq1)* strains showed almost no transcriptional changes compared to N2 in the absence of exogenous serotonin or loxapine. The lack of any transcriptional changes was especially striking for *ser-5(vq1)* with only 5 DEGs, one of them being *ser-5* itself (**supp Fig. 2**). At the transcriptional level, *ser-5(vq1)* has essentially no phenotype despite phenocopying the increased fat storage seen with loxapine treatment. The dramatic changes in transcription induced by loxapine treatment are, therefore, dispensable for the fat storage phenotype, suggesting that they are a secondary downstream response necessary to process incoming nutrients.

The transcriptional changes induced by either serotonin or loxapine treatment largely overlapped, with 70% of the loxapine response being a subset of the serotonin response whose expression was correlated by an R= 0.94. This transcriptional response was heavily enriched in DNA, RNA, Protein, and energy metabolism, further pointing to a response required to process the incoming nutrients. Consistent with that interpretation was that calorie restriction, an intervention where less food needs to be processed, caused transcriptional changes in the opposing direction (**Fig. 3**). Blocking the activity of the transcriptional responses, which included metabolism, mRNA function, and proteolysis dampened hyperphagia but also resulted in toxicity not allowing for a definitive interpretation (**Fig. 4**)

The experiments inactivating the germline by limiting translation provided the final piece of evidence that modulating fat storage thresholds can drive hyperphagia. Inactivating the germline by limiting translation-induced hyperphagia and increased fat storage while decreasing respiration (**Fig. 5**). Loxapine treatment still showed a slight increase in respiration that could be interpreted to contribute to the hyperphagic effect instead of the fat storage phenotype. In contrast, the inactivation of the germline reduced energy expenditure and induced hyperphagia, revealing that increased energy expenditure is not the only parameter that drives feeding^53^. The failure of loxapine or olanzapine to further increase feeding in germline-less animals is likely because the animals’ ability to store fat quickly reaches its maximum once the germline is inactivated (**Fig. 1G, H**, **Fig. 6C**). In contrast, serotonin still induces hyperphagia in animals with impaired germlines (**Fig. 5C**). Presumably serotonin is still able to induce hyperphagia because it promotes lipolysis (**Fig. 1G, H**) - and thus frees up the capacity to store fat^35,39,62–63^. As in *C. elegans*, removing the germline by ovariectomy in mice increased adiposity and blocked antipsychotic-induced metabolic side effects (**Fig. 6**). Taken together, these data show that fat storage and hyperphagia can be uncoupled and that it is possible to induce hyperphagia irrespective of fat storage.

Finally, it is noteworthy that antipsychotics are used to treat schizophrenia, a disease whose onset in young adulthood coincides with the start of the reproductive period in humans. Changes in lipid profiles predate the onset of psychosis, and lipid profiles of at-risk patients can be predictive of future psychosis ^64–66^. It is not clear if the metabolic effects of antipsychotics are only side effects or if they play a part in therapeutic efficacy. However, as shown previously, the metabolic side effects of antipsychotics are evolutionarily conserved between *C. elegans* and mice^8^, and we expect that the ability of the germline to modulate energy balance does influence antipsychotic-mediated metabolic side effects across all species.

## Methods

### *C. elegans* maintenance and strains

Worms were cultured on nematode growth medium (NGM) agar plates with *Escherichia coli* strain OP50 at 20 °C as described^67^. The N2 Bristol strain was obtained from the *Caenorhabditis* Genetic Center (CGC) and used as wild type in all assays. Worms All experiments were performed on either Day 1 or Day 4 adults. *C. elegans* studies did not require IRB or IACUC approvals at The Scripps Research Institute.

### C. elegans strains

N2, VV212 *ser-5(vq1),* VV222 *ser-7(vq2),* ESC332 *rpoa-2(cse319[degron::GFP::rpoa-2]) I; ieSi57 II.,* ESC351 *rpoa-2(cse319[degron::GFP::rpoa-2]) I; reSi2 II.,* ESC352 *qzIs15[rpoa-2p::degron::GFP::rpoa-2] I; ieSi60 II.,* ESC360 *rpoa-2(cse319[degron::GFP::rpoa-2]) I; emcSi71 IV.,* ESC373 *rpoa-2(cse319[degron::GFP::rpoa-2]) I; ieSi61 II.* ESC374 *rpoa-2(cse319[degron::GFP::rpoa-2]) I; ieSi38 IV*.

### Bacterial Clearance

Bacterial Clearance assay follows the protocol explained in depth in Clark & To *et al.* 2024^21^. L1 age synchronized worms were plated on 96 well microtiter plates at an initial volume of 120ul. On the L4 larval stage, FUDR was added to a final concentration of 0.12nM and a total volume of 150ul. On the D1 adult stage, the small molecule of choice was added to the plate, and the plates were shaken for 20 min on a plate shaker. Initial OD600 was taken at this time for bacterial clearance. On adult stage D4, plates were shaken again for 20 min on a plate shaker before the second OD600 reading was taken. Plates were kept if lifespan analysis was also being conducted. Unless being actively worked with, plates were kept in 20C incubators. Bacterial clearance analysis was completed using Excel and GraphPad Prism.

### Pharyngeal Pumping

L1 age synchronized worms were plated on 96 well microtiter plates at 120ul initial volume. On the L4 larval stage, FUDR was added to a final concentration of 0.12nM and a total volume of 150ul. On D1 adult stage, animals were dosed with small molecules of interest and shaken for 20 minutes on a plate shaker. Plates were incubated for six hours in 20C incubators. After six hours, the worms were transferred to NGM plates and allowed to equilibrate for 30 minutes. After 30 minutes, worms were tracked on a microscope for two minutes each, counting each time they participated in grinder movements. After two minutes, the worm was removed from the plate, and the next worm was tracked. Analysis was completed using Excel and GraphPad Prism.

### Oxygen Consumption Rates

L1 age synchronized worms were plated in on Seahorse e-flux plates (Agilent Technologies, Seahorse Bioscience, catalog number: 101085-004)) at a volume of 120ul at a concentration of 8-15 worms per well and with x-ray irradiated liquid culture (bacterial strain OP50 at 6mg/ml, S-complete medium with 50 µg ml−1 carbenicillin and 0.1 μg ml−1 fungizone). At the L4 stage, FUDR was added to a final concentration of 0.12nM and a total volume of 150ul. On Day 1, drug conditions were added with no more than 0.5% DMSO being added to each well if DMSO was needed to dissolve the drug of choice. Plates were shaken at 800rpm for 20min. Hydrate Sensor cartage at 37C with calibrant for at least 4 hours prior to assay but no more than 72h. On Day 4, plates were shaken for 20min at 800rpm prior to the assay. On the sensor plate, 17.2ul Carbonyl cyanide 4-(trifluoromethoxy) phenylhydrazone (FCCP) was added to injection port A (total final volume 10uM). In sensor plate injection port B, 19.4ul Sodium Azide was added (total final volume 40mM). Plates were then run through the assay with baseline having 5 readings with 30s mixing and FCCP and NaN3 readings having 4 readings with 30s mixing. Analysis was completed using Excel and GraphPad Prism. **Note:** The heater will not be on for the machine thus the results will vary based on the ambient temp, time of day, and buffer solutions. Control for these by performing the experiment at the same time of day.

### Oil Red O fat staining

Oil Red O staining was performed as described^39^. Worm populations were grown in accordance with the bacterial clearance assay described above^21^. After bacterial clearance phenotypes were confirmed for each drug on Day 4, worms were collected and washed using a 40uM cell strainer to strain and wash the D4 worms with phosphate-buffered saline (PBS) and incubated on ice for 10 min before fixation. Worms were then stained in filtered Oil Red O (Thermo Scientific) working solution (60% Oil Red O in isopropanol: 40% water) overnight, horizontally. Approximately 200 worms were fixed and stained for all genotypes within a single experiment. For each experimental condition, we observed around 100 worms on slides, after which 30-40 worms were randomly chosen for imaging. All experiments were repeated at least three times. Oil Red O-stained worms were imaged using 10× objective on a Zeiss Axio Imager microscope taken in DIC settings. Images were acquired with the Software AxioVision (Zeiss) and then analyzed using ImageJ software. Scale bars in all images indicate 20 μm.

### Lifespan

Lifespan assay follows the protocol explained in depth in Solis *et al.* 2011^68^. L1 age synchronized worms were plated on 96 well microtiter plates at a volume of 120ul. On the L4 larval stage, FUDR was added to a final concentration of 0.12nM and a total volume of 150ul. On the D1 adult stage, the small molecule of choice was added to the plate, and the plates were shaken for 20 min on a plate shaker. OD600 was taken at this time if bacterial clearance was also being measured. Worm populations were recorded for life and death every two to three days after the plates were shaken for 10 min on a plate shaker. Unless being actively worked with, plates were kept in 20C incubators. Lifespan analysis was completed using STATA and GraphPad prism.

### Length measurements

Worm morphological comparisons were imaged at 2X on a Nikon TMS inverted microscope. Worms were imaged in the liquid culture in 96 well plates as stated in the bacterial clearance method. IAA induction was started on L4, but images were taken on Day 4 of adult life or the terminal point of the bacterial clearance assay. Worm body length comparisons were made in ImageJ using a segmented line tool down the midline of each animal from head to tail. Real length comparisons were made by measuring the length on the 96 well plate and measuring the pixel length in ImageJ. A two-way ANOVA was used, as indicated in the figure legends.

### Auxin (IAA) treatment

The natural auxin IAA was purchased from Alfa Aesar (#A10556). Dry stocks were kept at −20°C in a light-protected box. At the L4 larval stage, IAA was dissolved in Dimethyl sulfoxide (DMSO) at 600X dilution factor; 0.25uM was added to each well in the 96 well plates before the plates were shaken at 800rpm on a microtiter plate shaker. Controls for experiments using IAA are wells that receive an equal amount of DMSO.

### RNA-seq

Worm populations were grown in accordance with the bacterial clearance assay described above. After bacterial clearance phenotypes were confirmed for each drug on Day 4, worms were collected and washed using a 40uM cell strainer to strain and wash the D4 worms. The worms were suspended in minimal PBS before being placed in a -80C freezer at least overnight or until extraction. Worms were broken open using a precellys tissue homogenizer and Trizol (Invitrogen cat# 15596-026) with Zirkonium and glass beads. A phenol-chloroform extraction was then completed before the samples were further cleaned using a Qiagen kit (Qiagen RNEasy Plus Micro Kit, CAT# 74034). RNA samples were then sent for sequencing at the Scripps Research Genomics Core Facility using an Illumina Nextseq2000. The sequencing read data was cleaned of contaminating adaptor sequences, and low-quality reads with ‘trimmomatic’ and quality tested with ‘fastqc’. Reads were then aligned to the *C. elegans* genome using ‘STAR’ and converted into a count matrix with ‘HTSeq-count’. Differential expression between groups was calculated and statistically tested by ‘DESeq2

### Statistics

The N2 (wild-type) animals were included as controls for every experiment. Fold change indicates change relative to untreated N2 unless otherwise indicated. Error bars represent the standard error of the mean (SEM). Students’ t-test and two-way ANOVA were used, as indicated in the figure legends. All statistical analyses were performed using GraphPad Prism 10 (GraphPad Software). Appropriate multiple comparison corrections were used for ANOVAs. \**p* < 0.05, \*\**p* < 0.01, \*\*\**p* < 0.001, *\*\*\*\*p*<0.00001 NS: not significant. For lifespan analysis, Survival data were analyzed by a Log-rank test using STATA.

### Mouse studies

All procedures are approved by the UCSD or TSRI IACUC committee. C57BL/6 female mice (stock #000664) were purchased from Jackson Labs at 9 weeks of age. All mice were fed normal chow until 10 weeks of age. Olanzapine was compounded into the high-fat diet (HFD) (45 kcal% from fat) at a concentration of 54 mg/kg of diet (OLZ-HFD, Research Diets, Inc., D09092903, New Brunswick, NJ) ovariectomy was conducted using method PMID: 38416648, and mice were allowed to recover for three weeks prior to antipsychotic treatment. Mice were fed OLZ-HFD for 4 weeks, and their food intake and body weight were measured twice per week.

## Acknowledgement

We want to thank Drs. Anabel Perez-Gomez, Greg Solis, and Rafael Gomez-Amaro for their pioneering work on this project, Drs Chung-Chih Liu and Supriya Srinivasan for their protocols and use of microscopes for Oil Red O staining, Dr. Shakib Omari for his microscope operations support, Alan To for his technical support, and all other members of the Petrascheck lab for their continuous input during this project. Seahorse experiments were completed thanks to the use of equipment from Dr. Enrique Saez’s laboratory. RNA-seq was completed thanks to Steve Head at The Scripps Research Institute (TSRI) Genomics Core Facility. CC was funded by the Dorris Neuroscience Scholar Fellowship. Some strains were provided by the Caenorhabditis Genetics Center CGC, funded by the NIH Office of Research Infrastructure Programs (P40 OD010440). NIH grants funded this project to M.P. R01 AG080376 and R01 DK117872.

---

**Supplemental to Figure 1:**
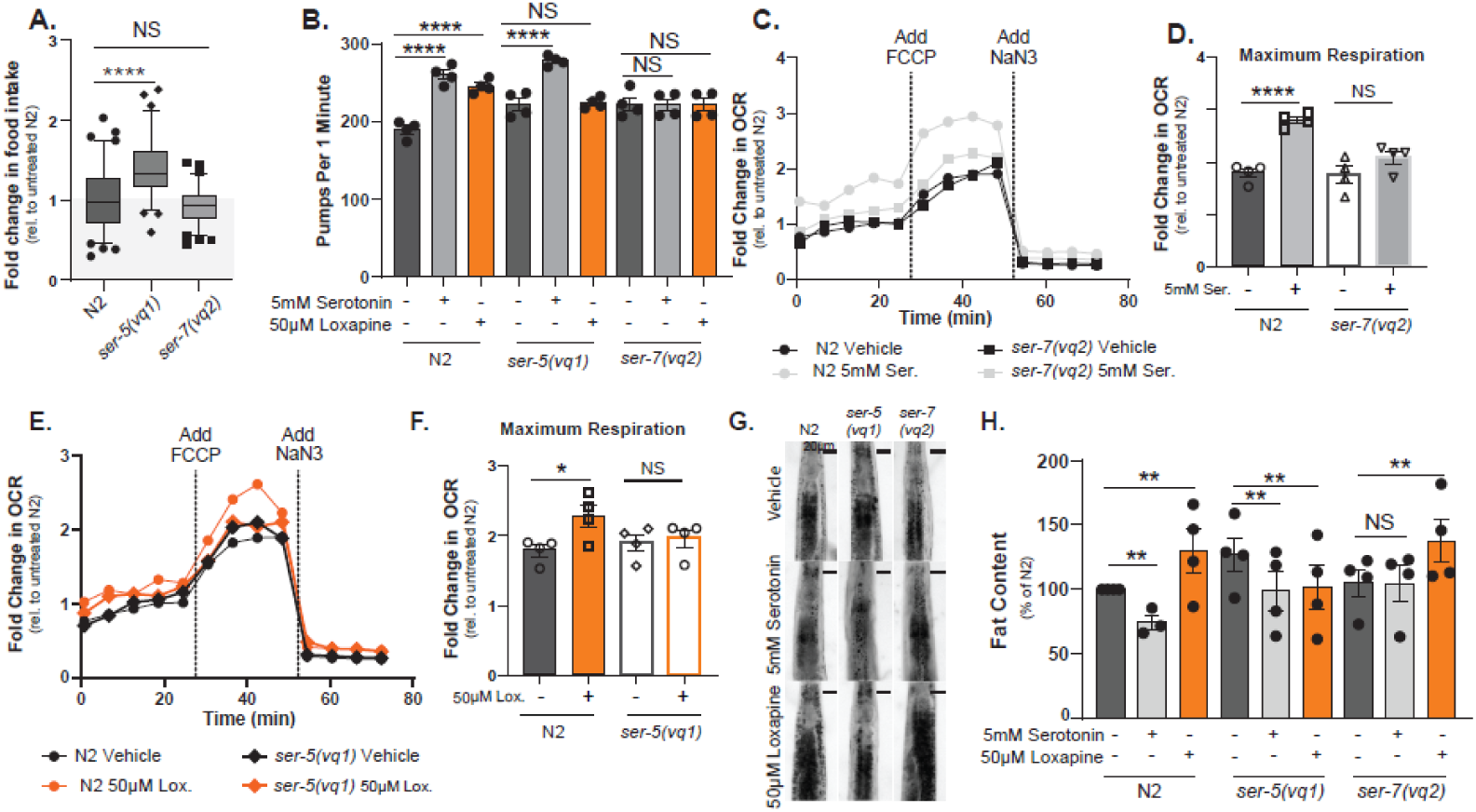
**(A).** Box blots representing the fold change in food intake of vehicle-treated N2 (wild-type), *ser-5(vq1)*, and *ser-7(vq2)* mutants. **(B)** Bar graphs depicting mean pharyngeal pumping rates of day 1 adults treated with vehicle, serotonin (5mM), or loxapine (50μM) for six hours. Each dot represents a trial of n=20 animals each. (**C**) Fold change in oxygen consumption rate (OCR) relative to control measured for vehicle-treated N2 (wild-type) and *ser-7(vq2)* mutants treated with vehicle (black) or 5mM Serotonin (light grey) for 72h measured on a Seahorse Extracellular Flux (XF) analyzer. **(D)** Bars show the maximum respiration rate measured for vehicle-treated N2 (wild-type), *ser-*and *ser-7(vq2)* mutants treated with vehicle (black) or 5mM Serotonin (light grey). Each dot represents a trial of n=100 animals each. (**E**) Fold change in oxygen consumption rate (OCR) relative to control measured for vehicle-treated N2 (wild-type), *ser-*and *ser-5(vq1)* mutants treated with vehicle (black) or 50μM loxapine (orange) for 72h measured on a Seahorse Extracellular Flux (XF) analyzer. **(F)** Bars show the maximum respiration rate measured for vehicle-treated N2 (wild-type) and *ser-5(vq1)* mutants treated with vehicle (black) or 50μM loxapine (orange). Each dot represents a trial of n=100 animals each. **(G)** Representative images of vehicle, serotonin, or loxapine-treated N2 (wild-type), *ser-5(vq1)*, and *ser-7(vq2)* mutants stained with Oil Red O to visualize fat depots. **(H)** Quantification of fat staining of day 4 adult N2, *ser-5(vq1)*, or *ser-7(vq2)* mutants treated with either vehicle (dark grey), serotonin (light grey), or loxapine (orange). Bars show the mean % fat content relative to vehicle-treated N2 controls of 4 trials consisting of n=20 animals each. Asterisks indicate significance, \*\**p* < 0.05, \*\**p* < 0.01, \*\*\**p* < 0.001, *\*\*\*\*p*<0.00001 NS: not significant Abbreviation Ser. for serotonin and lox. for loxapine.

**Supplemental for Figure 2:**
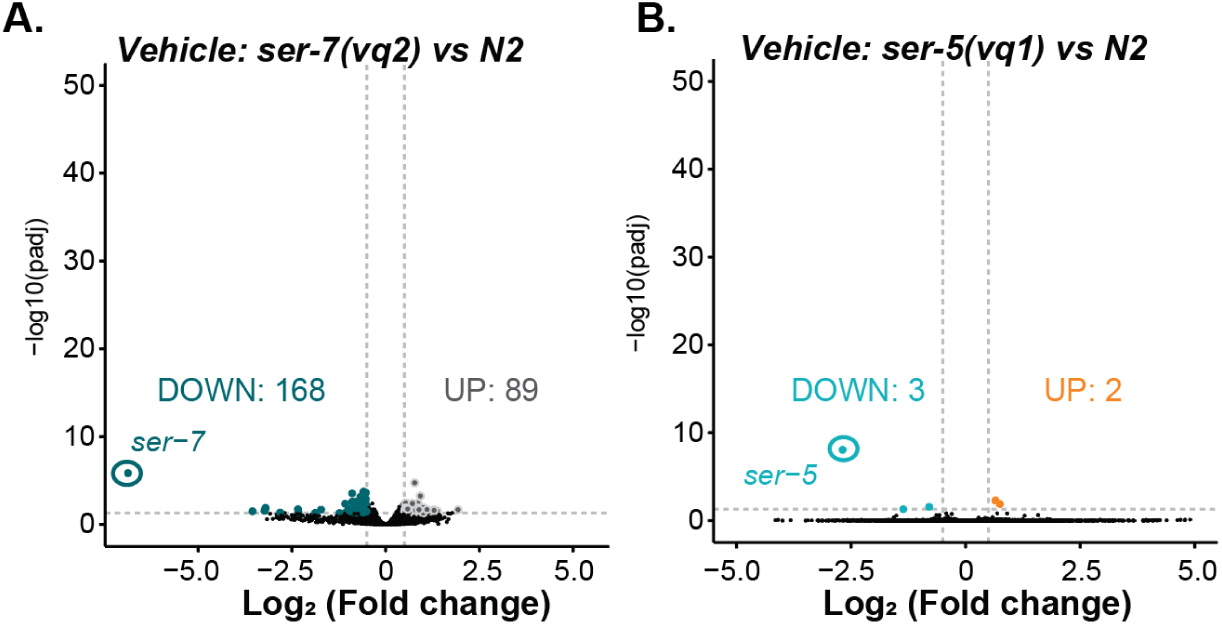
**Title** Graphs A-B show volcano plots representing -Log_10_ (pvalues) as a function of Log_2_(Fold changes) in gene expression for 14,912 genes detected in all samples. Differentially expressed genes (DEGs) are colored. **(A).** Gene expression changes in *ser-7(vq2)* mutants compared to N2 animals. **(B)** Gene expression changes in *ser-5(vq1)* mutants compared to N2 animals. In the absence of an exogenous ligand, gene-expression profiles of *ser-5(vq1)* and *ser-7(vq2)* mutants are very similar to those of N2 animals. For all volcano plots, FDR ≥ 0.05.

**Supplemental for Figure 4:**
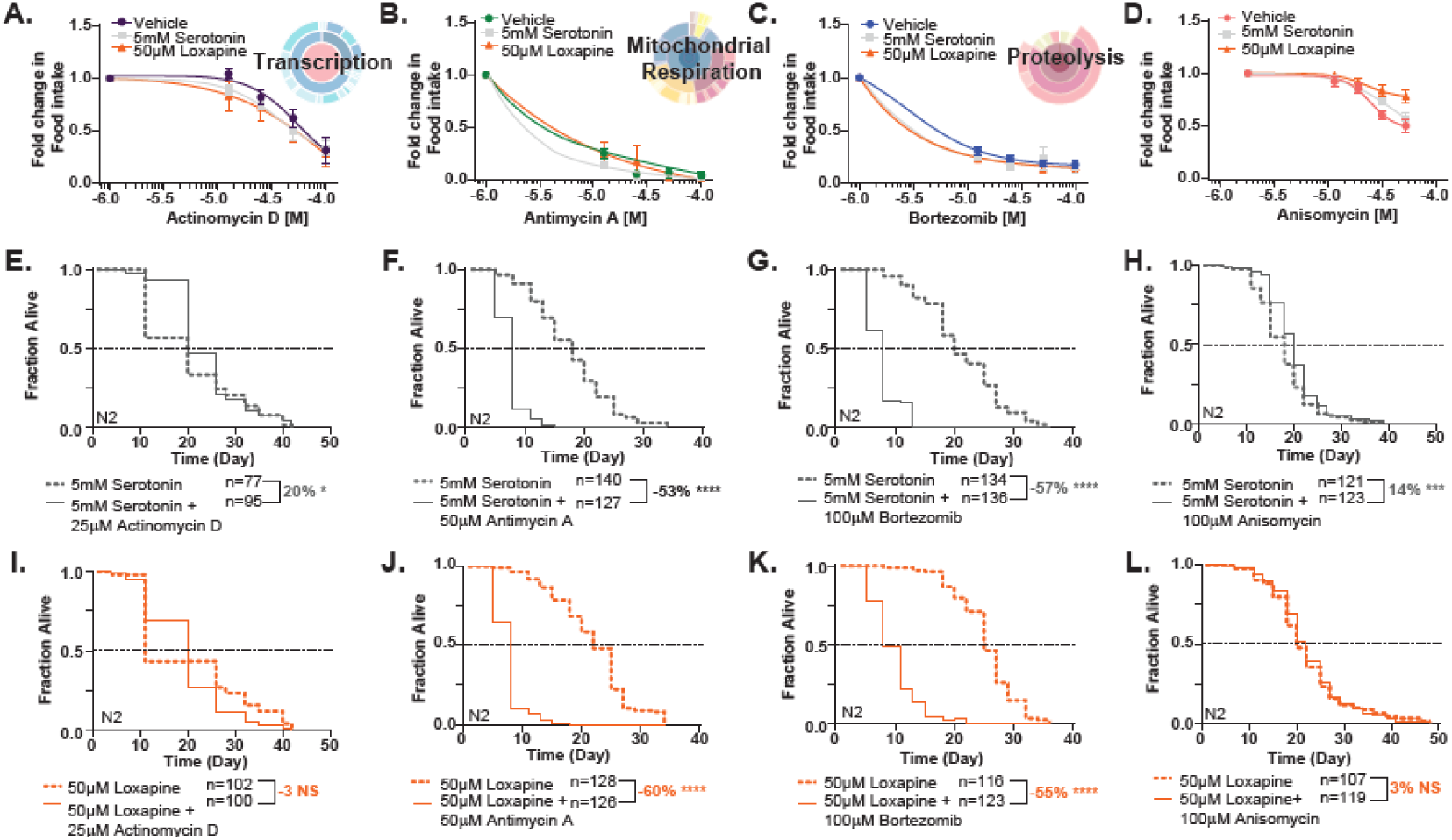
The graphs A-D show the average fold change in food intake similar to what is shown in actual Fig. 4(A-D). However, the actual figure 4 in the text normalized all values relative to the N2 vehicle control values, while the supplemental figure 4 normalizes the values for each curve relative to the no inhibitor control values. **(A)** Dose-response curve depicting the fold change in feeding as a function of increasing doses of the transcription inhibitor Actinomycin D. **(B)** Same as (A) but for the transcription inhibitor mitochondrial respiration inhibitor Antimycin A. **(C)** Same as (A) but for the proteasome inhibitor bortezomib. **(D)** Same as (A) but for the translation inhibitor anisomycin. Graphs **E-H** show Kaplan-Meier survival curves for N2 animals co-treated with serotonin (5mM) and the same inhibitors as in A-D. **(E)** Kaplan-Meier survival curves for N2 treated with Actinomycin D and serotonin. **(F)** Kaplan-Meier survival curves for N2 treated with Antimycin A and serotonin. **(G)** Kaplan-Meier survival curves for N2 treated with bortezomib and serotonin. **(H)** Kaplan-Meier survival curves for N2 treated with anisomycin and serotonin. Graphs **I-L** show Kaplan-Meier survival curves for N2 animals co-treated with serotonin (5mM) and the same inhibitors as in A-D. **(I)** Kaplan-Meier survival curves for N2 treated with bortezomib and loxapine. **(J)** Kaplan-Meier survival curves for N2 treated with Antimycin A and loxapine. **(K)** Kaplan-Meier survival curves for N2 treated with Antimycin A and loxapine. **(L)** Kaplan-Meier survival curves for N2 treated with anisomycin and loxapine. Survival data were analyzed by a Log-rank test. Asterisks indicate significance, \*\**p* < 0.05, \*\**p* < 0.01, \*\*\**p* < 0.001, *\*\*\*\*p*<0.00001 NS: not significant Abbreviation Ser. for serotonin and lox. for loxapine.

